# GPNMB and glycosphingolipid measurements in cerebrospinal fluid and plasma from Parkinson’s disease patients

**DOI:** 10.64898/2026.04.09.712000

**Authors:** Maria E Fernandez-Suarez, Reuben Bush, Erika Di Biase, Danielle te Vruchte, David A Priestman, Mario Cortina-Borja, Oliver Cooper, John Hardy, Penelope J Hallett, Ole Isacson, Frances M Platt

**Affiliations:** Department of Pharmacology, University of Oxford, Oxford, OX1 3QT, UK; Aligning Science Across Parkinson’s (ASAP) Collaborative Research Network, Chevy Chase, MD, USA; Department of Biochemistry, University of Oxford, Oxford, OX1 3QU, UK; Neuroregeneration Institute, McLean Hospital / Harvard Medical School, Belmont, MA, 02478, USA; Population, Policy, and Practice Research and Teaching Department, UCL Great Ormond Street Institute of Child Health, University College London, London, UK; Department of Neurodegenerative Disease, UCL Institute of Neurology, UK; UK Dementia Research Institute at UCL, London, UK

## Abstract

**Background:** Parkinson’s disease (PD) is a prevalent neurodegenerative disorder characterized by progressive motor dysfunction and broad cellular impairment, including significant disruptions in lysosomal function, lipid metabolism, and intracellular trafficking. Glycosphingolipids (GSLs), critical for various cellular processes, depend on effective lysosomal degradation. Aberrant GSL metabolism has been linked to PD pathology, and glycoprotein non-metastatic melanoma protein B (GPNMB) has emerged as a biomarker associated with lysosomal dysfunction and lipid imbalance in PD

**Objectives:** To assess the relationship between GPNMB and GSL levels in cerebrospinal fluid (CSF) and plasma from PD patients and controls within the BioFIND cohort. We also investigated potential sex differences and associations with PD-related biomarkers such as α-synuclein

**Methods:** GSL species and GPNMB protein levels were quantified using high-performance liquid chromatography (HPLC) and ELISA assays, respectively, in matched CSF and plasma samples from PD patients and controls

**Results:** Levels of the paraglobosides GSL species, alpha-2,3SpG and pGb were significantly elevated in the plasma of PD patients compared to healthy controls, while levels of the ganglioside GD1a and the lacto-series GSL, Le^b^ combined (GD1a + Le^b^), were significantly reduced in PD. GPNMB levels positively correlated with several GSL species in both plasma and CSF. Plasma GSLs and GPNMB concentrations were significantly higher in females compared to males, independent of PD diagnosis. CSF GPNMB correlated positively with age and α-synuclein concentrations

**Interpretation:** Our findings confirm that GSL metabolism is altered in PD. They also highlight significant sex-based biochemical variations in GSL and GPNMB levels, emphasizing the need for sex-specific analyses in PD biomarker research. The relationship between GSLs and GPNMB supports their potential as interconnected biomarkers of lipid pathology in PD.

## Introduction

Parkinson’s disease (PD), the second most common neurodegenerative disorder, affects over 10 million people and is marked by progressive motor decline and cellular dysfunction. Early disruptions in lysosomal function, lipid metabolism, and intracellular trafficking are increasingly recognized as central to PD pathogenesis, impacting neurons and glia and indicating systemic failure of proteostatic and lipid regulation ^1–4^. While most cases are idiopathic, genetic variants, especially heterozygous mutations in lysosomal storage disorder (LSD) genes like *GBA1* are enriched among PD risk alleles, pointing to lysosomal dysfunction in both familial and sporadic forms. Moreover, established environmental and pharmacological PD risk factors, including paraquat and MPP+, have also been shown to impair lysosomal function, reinforcing the hypothesis that compromised autophagy–lysosomal degradation pathways predispose to disease ^5, 6^.

The lysosome, the cell’s main catabolic compartment and metabolic signaling hub, clears misfolded proteins and degrades lipids. Its dysfunction promotes accumulation of neurotoxic aggregates like α-synuclein, contributing to neuronal vulnerability. Aging, the strongest risk factor for PD, is linked to reduced lysosomal efficiency and altered lipid profiles, highlighting lysosomal failure as a key mechanism in age-related neurodegeneration ^7–8^. Glycosphingolipids (GSLs), which rely on lysosomal degradation, play roles in membrane structure, signaling, trafficking, and immunity. Defects in GSL-degrading enzymes cause substrate build-up typical of LSDs. Mutations affecting GSL catabolism are associated with PD risk, linking GSL metabolism, lysosomal function, and neurodegeneration.

Our previous work identified reductions in multiple lysosomal glycosidases in the substantia nigra of PD patients, alongside a marked loss of complex gangliosides, changes exacerbated by aging ^8–10^. Alterations in GSL profiles were also detected in plasma and cerebrospinal fluid (CSF), even at prodromal stages of disease, as indicated by individuals with REM sleep behavior disorder, underscoring the potential of GSLs as biomarkers of early pathology ^8^. One lysosome-associated molecule emerging at the intersection of lipid metabolism and neuroinflammation is glycoprotein non-metastatic melanoma protein B (GPNMB). GPNMB is a transmembrane glycoprotein upregulated in lysosomal storage conditions and modulated by inflammatory cues ^11–16^ and has been identified as a PD risk locus in genome-wide association studies ^17^, with elevated mRNA expression in the PD substantia nigra ^18^. Recent findings demonstrate that GPNMB localizes to the lysosome during stress and is secreted via lysosomal exocytosis in a LRRK2-dependent manner ^19^, positioning it as a key readout of lysosomal stress and lipid perturbation. In experimental systems, pharmacological inhibition of GBA1 induces robust GPNMB expression, and elevated levels of GPNMB protein have been observed in PD brain, correlating with lipid accumulation and disease severity ^9, 20^.

Despite these findings, previous studies examining GSL and GPNMB changes in PD were limited as they used independent cohorts for CSF and plasma analyses and included a preponderance of male participants, restricting generalizability and hindering the assessment of sex-specific differences. To address these limitations, the current study leveraged the BioFIND cohort, with paired CSF and plasma samples from male and female PD patients and healthy controls. This allowed us to directly examine the relationship between GSLs and GPNMB across biofluids and assess their association with clinical parameters, including α-synuclein levels, age, and sex.

## Materials and Methods

The concentration of different species of glycosphingolipids (GSLs) was measured in the plasma and CSF of 204 and 157 subjects, respectively, belonging to the BioFIND study cohort ^21^, the BioFIND institutional review board approved the study protocol and informed consent was obtained from each participant. There were six *GBA1* carriers in the plasma samples analyzed (for GSLs and GPMNB) and five and seven in CSF for GSLs and GPNMB, respectively. The original GSL qualitative method we developed was published in 2004 ^22^ and measured the oligosaccharides selectively released from glycosphingolipids using a ceramide glycanase enzyme derived from the medicinal leech. The protocol was adapted to achieve sensitive and reproducible quantitation of GSLs in control and patient plasma ^23^ and CSF samples ^24^. The method uses the fluorescent compound anthranilic acid (2-AA) to label oligosaccharides prior to analysis using normal-phase high-performance liquid chromatography. With the inclusion of a 2AA-labelled chitotriose calibration standard (Ludger Ltd., United Kingdom), it is possible to obtain molar quantities of individual GSL species. Detailed protocols for plasma and CSF processing are available at protocols.IO ^23, 24^ and the BioFIND Data Repository at Laboratory of Neuro Imaging (LONI) (https://biofind.loni.usc.edu/download-data.php). After quantification, to compare between sample sets, all the values were normalized using the total GSLs concentration from a Pool sample present in all sample sets and using the following equation for plasma:

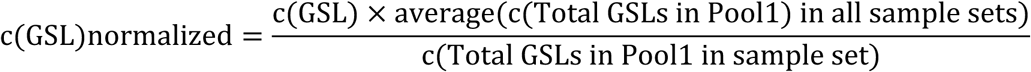

and for CSF:

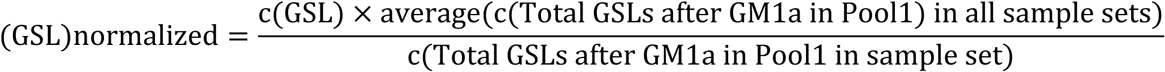

Because CSF samples were run in duplicate the final values were obtained by calculating the mean across the two runs. In plasma samples, due to close co-elution of two pairs of peaks: GD1a + Le^b^ and GD1b + GD1-alpha, the individual GSL species in these pairs could not be resolved and have therefore been reported as the sums of the respective combined peaks. GPNMB protein concentration was measured in the plasma of 205 subjects and in the CSF of 185 subjects belonging to the BioFIND study cohort. Every sample was identified with a bar code and the evaluation of protein concentration was performed blindly by the colorimetric ELISA using a commercial kit (ELH-Osteoactivin, RayBiotech) (Moloney et al 2018).

The plasma samples were diluted 50 times into the diluent buffer A (Item D). 50 times dilution provided the absorbance values falling into the linear range of the standard curve in pilot optimizing assays. Calibrator peptide (Item C) dilutions were prepared in the same buffer of the samples. Ten 96-multi-well plates were used to run all the samples in triplicate following the manufacturer’s instructions. In brief, 100 μl of each standard and sample were added to the appropriate wells of a coated 96 multi-well plate (Item A) and the plate was incubated for 2.5 hours at room temperature with gentle shaking. The wells were washed 4 times with 1X wash solution (Item B) and 100 μl of biotinylated antibody (diluted 80-fold with 1X dilution buffer B) was added. The plate was incubated 1 hour at room temperature with gentle shaking. The wells were rinsed 4 times in 1X wash buffer and 100 μl of HRP-Streptavidin solution (Item G, 500-fold diluted in 1X dilution buffer B) was added to each well. The plate was incubated for 45 minutes at room temperature with gentle shaking. The wells were rinsed 4 times with 1X wash buffer and 100 μl of TMB One-Step Substrate Reagent (Item H) was added to each well. The plate was incubated 30 minutes at room temperature in the dark with gentle shaking. At the end of the incubation 50 μl of Stop Solution (Item I) was added to each well and the absorbance at 450 nm was immediately read with the SPECTRAmax plate reader (Molecular Devices). Plasma and CSF sample pools, one deriving from the control group and one deriving from the PD group, were assayed in each plate together with sample-tests and standards, following the dilution in the diluent buffer. Values deriving from one pool were used for normalization purposes.

To obtain the GPNMB concentration in the plasma and CSF samples, normalized over the pool, the absorbance values were processed as follows: the protein concentration of the pools in each plate was interpolated by the standard curve and expressed as ng/ml. Per each plate, the GPNMB values of the samples were expressed as ELISA Unit by dividing the sample absorbance value by the average of the pool absorbance values, following the equation (1):

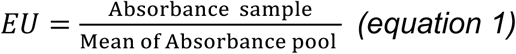

To obtain the GPNMB concentration of each sample test, the EU was multiplied by the average of the pool’s protein concentrations across the plates and by the dilution factor (50), following the equation in (2):

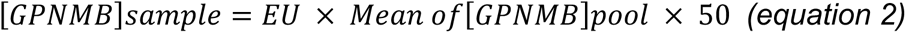

### Blinding

All authors were blind to the experimental groups during the experimentation, data acquisition and processing.

### Data availability and statistical analysis

All data were downloaded from the BioFIND website (https://biofind.loni.usc.edu/download-data.php) on October 28^th^, 2024. All study subjects were either cognitively intact or exhibited mild cognitive impairment (MCI). MCI was defined as subjects whose Montreal Cognitive Assessment (MOCA) score was < 26. Statistical analyses were performed using Prism version 10, GraphPad, Dotmatics, Boston, USA, and R v4.3.0, R Foundation for Statistical Computing, Vienna Austria.

## Results

### GPNMB protein levels are positively correlated with GSLs in the CSF and plasma

Given our previous findings of increased levels of GPNMB and GSLs in the substantia nigra in Parkinson’s disease (PD) ^8–10, 20^, this study evaluated GPNMB and GSL levels in cerebrospinal fluid (CSF) and plasma biofluids from subjects with PD and age-matched controls from the BioFIND cohort. Demographics and clinical characteristics of BioFIND participants have been reported previously ^25^. As outlined in Supplemental Table 1, a total of 157 and 185 CSF samples were analyzed for GSLs (92 PD, 65 controls) and GPNMB (107 PD, 78 controls), respectively. Within CSF samples analyzed for GSLs, the number of samples from males and females was 31 male and 34 female controls, and 54 male and 38 female PD patients. Within the CSF samples analyzed for GPNMB the number of samples from males and females was 36 control males and 42 control females, and 64 male and 43 female PD patients. In plasma samples, a total of 204 samples from the BioFIND cohort were analyzed for GSLs (116 PD subjects and 88 age-matched controls), and 205 for GPNMB (116 PD subjects and 89 age-matched controls). All samples analyzed for GSLs were also analyzed for GPNMB. In plasma, the sample number from male and females for analysis of GSLs was 44 female and 44 male controls, and 44 female and 72 male PD, and for analysis of GPNMB was 44 female and 45 male controls, and 44 female and 72 male PD. Concentrations of GPNMB protein and GSLs broadly differed between the subjects’ CSF and plasma samples. The concentration of GPNMB protein was 7-fold higher in plasma than CSF (plasma 23.74 ± 3.15 ng/ml, CSF 3.25 ± 0.5 ng/ml). Mean concentrations of GSLs in plasma ranged between 0.012 – 4.25 nmol/ml and CSF ranged between 3.26 – 26.16 pmol/ml. The high plasma-to-CSF GPNMB ratio likely reflects greater peripheral production by monocyte-macrophage lineages, contrasted with more limited production primarily in glial cells in the CNS. In addition, GPNMB is a large glycosylated protein that does not easily cross the blood-brain or blood-CSF barriers. As a result, absolute GPNMB concentrations are markedly lower in CSF than plasma.

GPNMB protein levels in the CSF were significantly correlated with all GSL species measured (**Figure 1A**). GSL biosynthesis is complex (**Supplementary Figure 1**) and the relative abundance of the different GSLs is unique for each tissue or cell type. When analyzed by HPLC, GSLs therefore generate distinctive profiles depending on the type of sample analyzed. The CSF HPLC profile (**Supplementary Figure 2**) is dominated by gangliosides and contains prominent peaks of GM1a, GD1a, GD1b and GT1b (a-series and b-series), and smaller peaks of GM3 and GD3 (precursors of a-series and b-series), which correspond to the major GSLs expressed in the mammalian brain. The strongest correlations of GPNMB with GSLs corresponded to the complex gangliosides GD3 and GD1a. There was also a significant positive correlation between GPNMB and total GSLs in the CSF. Individual GSL species were also correlated with each other, irrespective of which individual species were compared, with the strongest correlations between each species and their precursors and derivatives in the biosynthetic pathway (**Supplementary Figure 1)**. For example, GT1b levels were strongly correlated with its derivative GQ1b and its immediate precursors GD1b and GD1a, but to a lesser extent with the more distant GM3.

**Figure 1:**
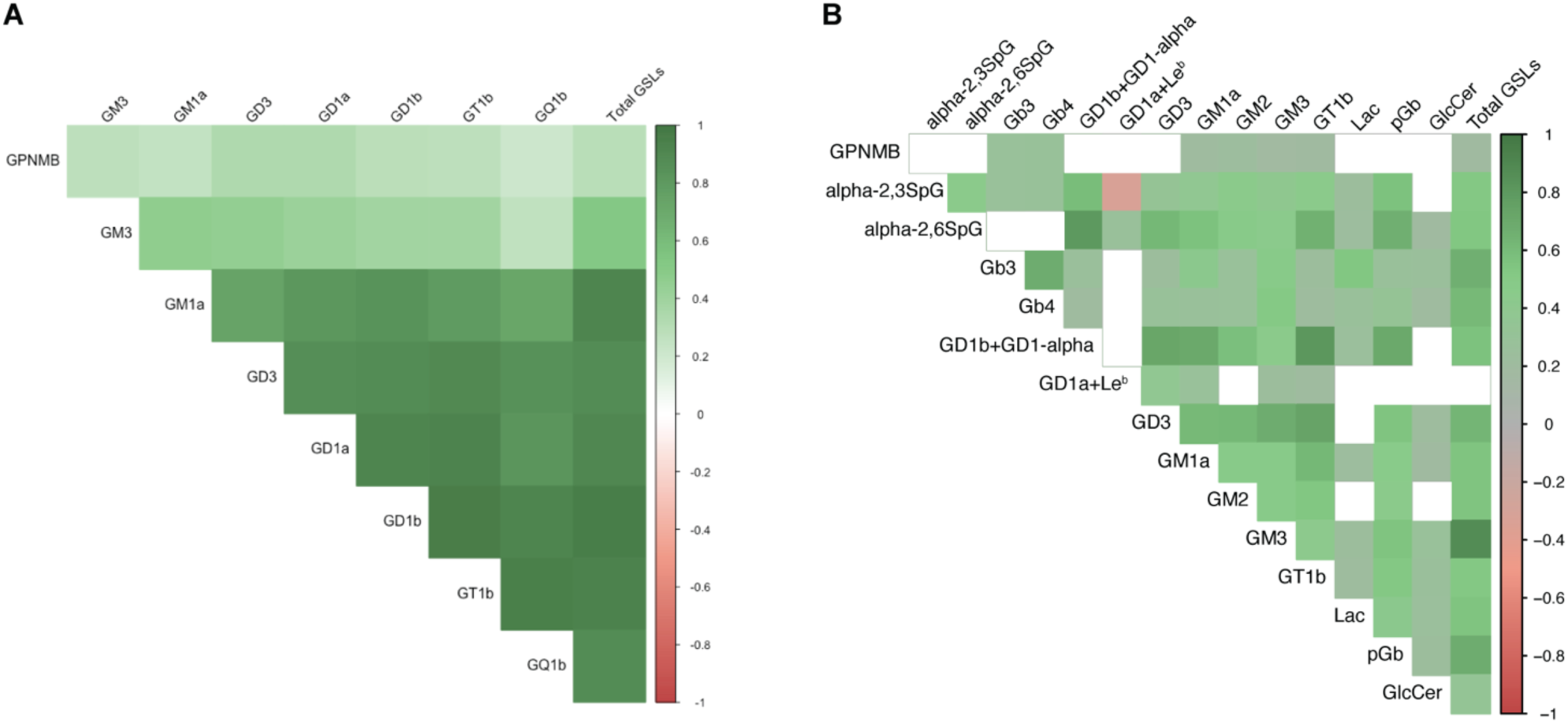
GSLs are highly correlated within subjects and positively correlated with GPNMB levels, in CSF and plasma. Spearman-rank correlations of CSF GSLs and GPNMB in CSF (A) and plasma (B), visualized as correlograms. Shading of cells represents the magnitude of correlation coefficient, statistically non-significant correlations visualized as blank cells. Threshold for statistical significance was *p* < 0.01.

As in the CSF, plasma GPNMB and GSL levels were significantly positively correlated (**Figure 1B**). The human plasma GSL profile contains approximately 14 peaks corresponding to 16 different GSL species including globo-series GSLs that are absent from brain and CSF (**Supplementary Figures 2 and 3**). We systematically analyzed 16 of these species. Very minor peaks without definitive structural assignment were not included in the analysis but were included in determining total GSL. In plasma, peaks corresponding to the species GD1a and Le^b^ could not be resolved as they closely co-elute. Hence, they were reported together as GD1a + Le^b^. Similarly, levels corresponding to the species GD1b and GD1-alpha were also reported together. In contrast to CSF, plasma contains fewer gangliosides (predominantly GM3) and contains more neutral GSL species. A representative profile is shown in **Supplementary Figure 3**. GPNMB was significantly correlated with globosides Gb3 and Gb4 and with gangliosides GD1a, GD1b, GM1a, GM2, GM3 and GT1b as well as with total GSLs (**Figure 1B**). There was a positive correlation between most GSL species when measured in the plasma. However, GD1a + Le^b^ and alpha-2,3SpG were negatively correlated with each other (**Figure 1B**). Interestingly, there was no correlation between any GSL species when plasma was compared with matched CSF samples (**Supplementary Figure 4**).

### Plasma levels of the GSLs alpha-2,3SpG, pGb, and GD1a + Le^b^ are significantly changed in PD

In plasma, from the 14 GSL species analyzed, three were statistical significantly different in PD compared to control subjects (**Figure 2**). We found that alpha-2,3SpG and pGb (a metabolic immediate precursor of alpha-2,3SpG) were significantly increased in PD whereas GD1a + Le^b^ was significantly reduced in the plasma of PD subjects compared to controls (**Figure 2**). No significant differences were found in plasma levels of GPNMB between PD and controls. Similarly, in the CSF, no significant differences were observed between PD and healthy controls in GPNMB or any of the GSLs assessed. Interestingly, when plasma GSLs were stratified by sex we found a significant decrease in males compared to females in alpha-2,3SpG, pGb, and GD1a + Le^b^ that was independent of disease status (**Figure 2**). This led us to further investigate the effect of sex on the levels of GSL species that were not changed in PD.

**Figure 2:**
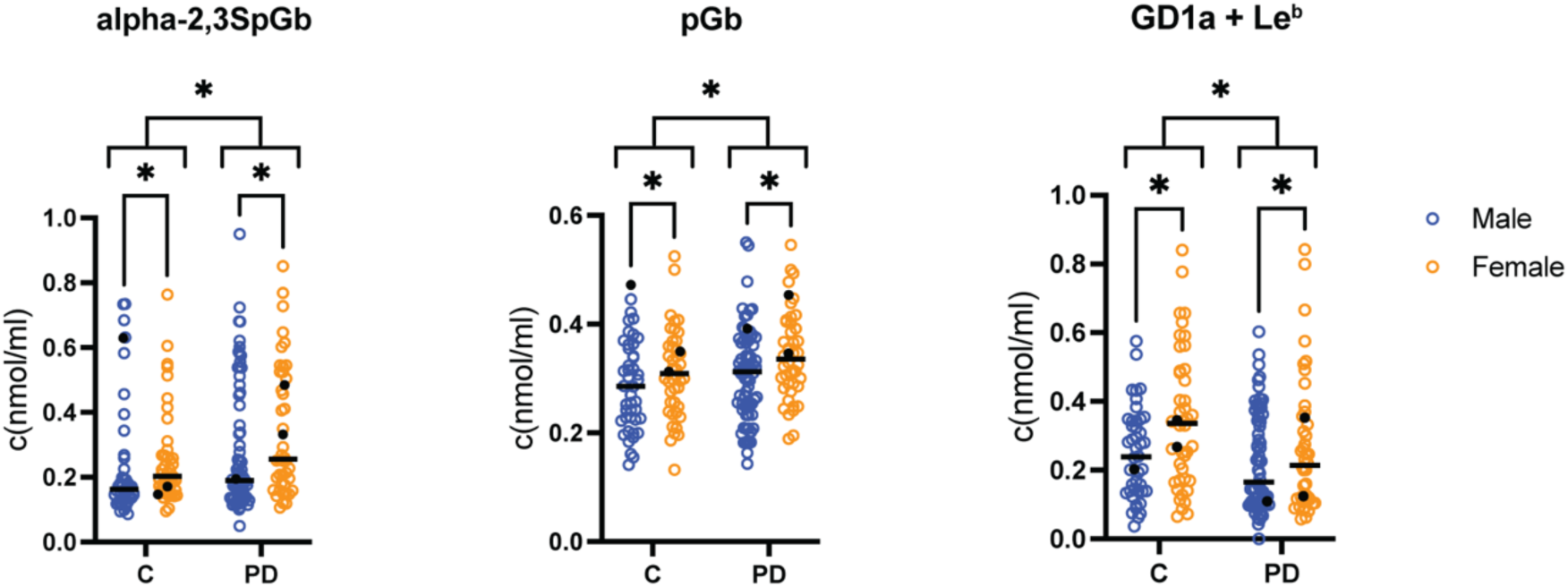
Levels of GSLs are significantly altered in the plasma of PD patients compared to controls. Quantified individual GSL species from plasma (*n* = 204), stratified by disease status (*n* = 88 controls and 116 PD) and sex (*n* = 88 female and 116 male). Individual subjects are visualized as circles, with controls (blue), PD (orange), and *GBA1* mutant carriers as filled in black circles (*n* = 6). The median for each group is indicated with a black line. Linear regression of log-normalized values of plasma GSL species, fitting variables by disease status and sex, \**p* < 0.05.

### GSL and GPNMB levels are decreased in the plasma of males whereas GPNMB is increased in males CSF

Given the reported 1.5-fold increased incidence rate of PD in males versus females ^26^, we explored the effect of sex on GSLs and GPNMB levels in plasma and CSF from control and PD individuals. In the plasma, 9 of the 11 GSL species not affected by disease status, as well as total gangliosides and total GSLs were found to be significantly higher in female subjects compared to males (**Figure 3).** This included the paragloboside alpha-2,6SpG and the globosides Gb3 and Gb4. The gangliosides, GD3 and GT1b from the b-series and GM1a, GM2 and GM3 from the a-series were also higher in females compared to male subjects (**Figure 3 and Supplementary Figure 1**).

**Figure 3:**
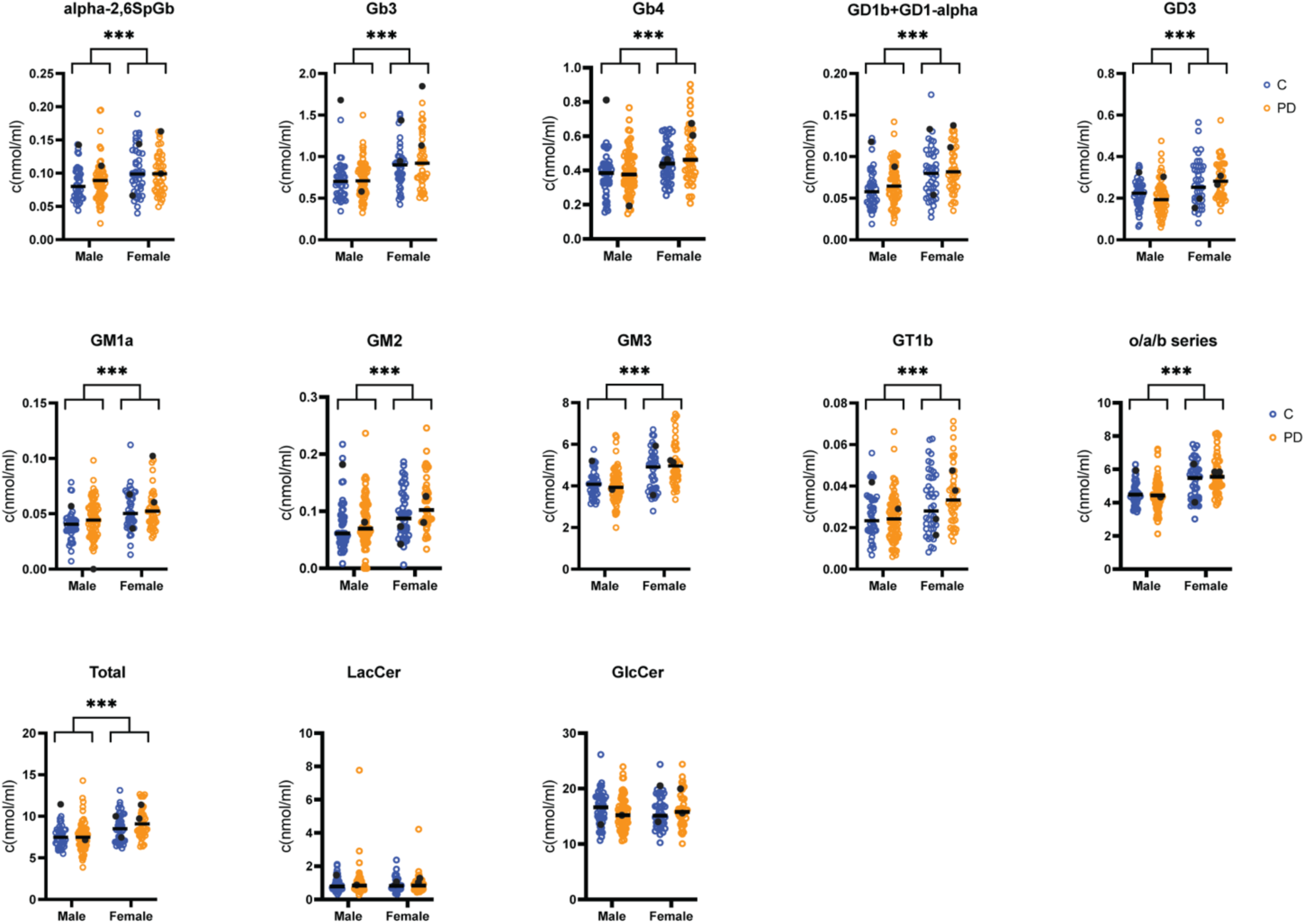
GSLs in plasma are decreased in male subjects compared to females irrespective of disease status. Quantified individual GSL species from plasma (*n* = 204), stratified by sex (*n* = 88 female and 116 male) and disease status (*n* = 88 controls and 116 PD). Individual subjects are visualized as circles, with controls (blue), PD (orange), and *GBA1* mutant carriers as filled black circles (*n* = 6). The median for each group is indicated with a black line. The o/a/b series do not include GD1a as it cannot be completely separated from Le^b^. One female PD subject was excluded from the analysis due to an abnormally high LacCer level (15 SD higher than mean) that therefore affected the total GSL level. Therefore, *n* = 203 for total GSLs allowing after the removal of this sample. Linear regression of log-normalized values of plasma GSL species, fitting variables by sex and disease status, *** *p* < 0.001.

We found no differences by sex or disease status in LacCer or GlcCer plasma levels. GPNMB protein levels in plasma were also statistically significantly higher in females than in males (**Figure 4A**). Conversely, in CSF GPNMB protein levels were significantly increased in males compared to females (**Figure 4B**). No significant differences were observed between males and females in any of the GSLs assessed in the CSF (**Supplementary Figure 5**). However, there was an overall trend towards lower levels of CSF GSLs in male PD subjects when compared to male controls (**Supplementary Figure 5**).

**Figure 4:**
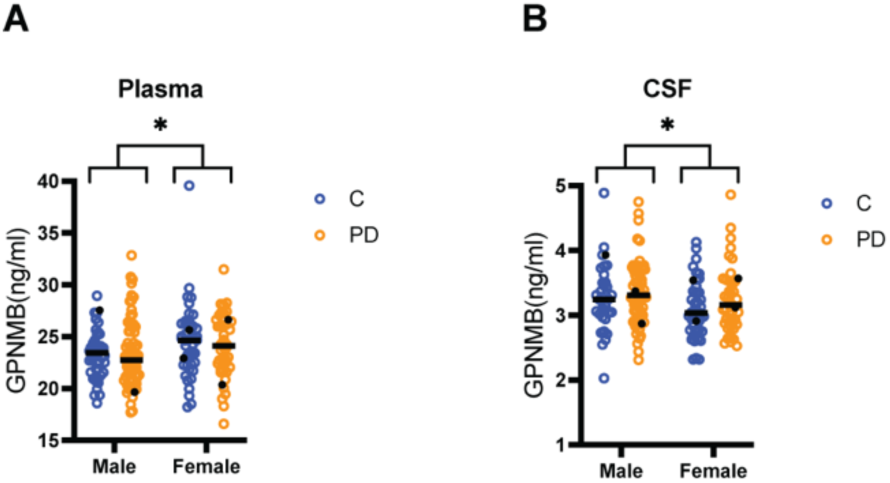
GPNMB levels are decreased in the plasma and increased in the CSF of male subjects compared to females. (A) Plasma GPNMB protein levels stratified by sex (*n* = 88 female and 117 male) and disease status (*n* = 89 controls and 116 PD). (B) CSF GPNMB protein levels stratified by sex (*n* = 85 female and 100 male) and disease status (*n* = 78 controls and 107 PD). Individual subjects are visualized as circles, with controls (blue), PD (orange), and *GBA1* mutant carriers as filled in black circles (*n* = 6 for plasma and *n* = 7 for CSF). The median for each group is indicated with a black line. Males vs Females were compared by Wilcoxon-Mann-Whitney test \**p* < 0.05.

Next, the relationship between GPNMB and GSL levels in the CSF was explored by sex. When stratifying by sex and disease status (**Table 1**), we found that the GSL and GPNMB correlation was driven by the female cohort. In female control and PD subjects, there was a statistically significant correlation between GPNMB and GD1a, GD1b, GD3, GM3, GQ1b and GT1b. Interestingly, this correlation was statistically significant only in PD females for GM1a and GM1b. There was no significant correlation in male subjects between any of the GSLs measured and GPNMB levels in the PD or control groups.

**Table 1:**
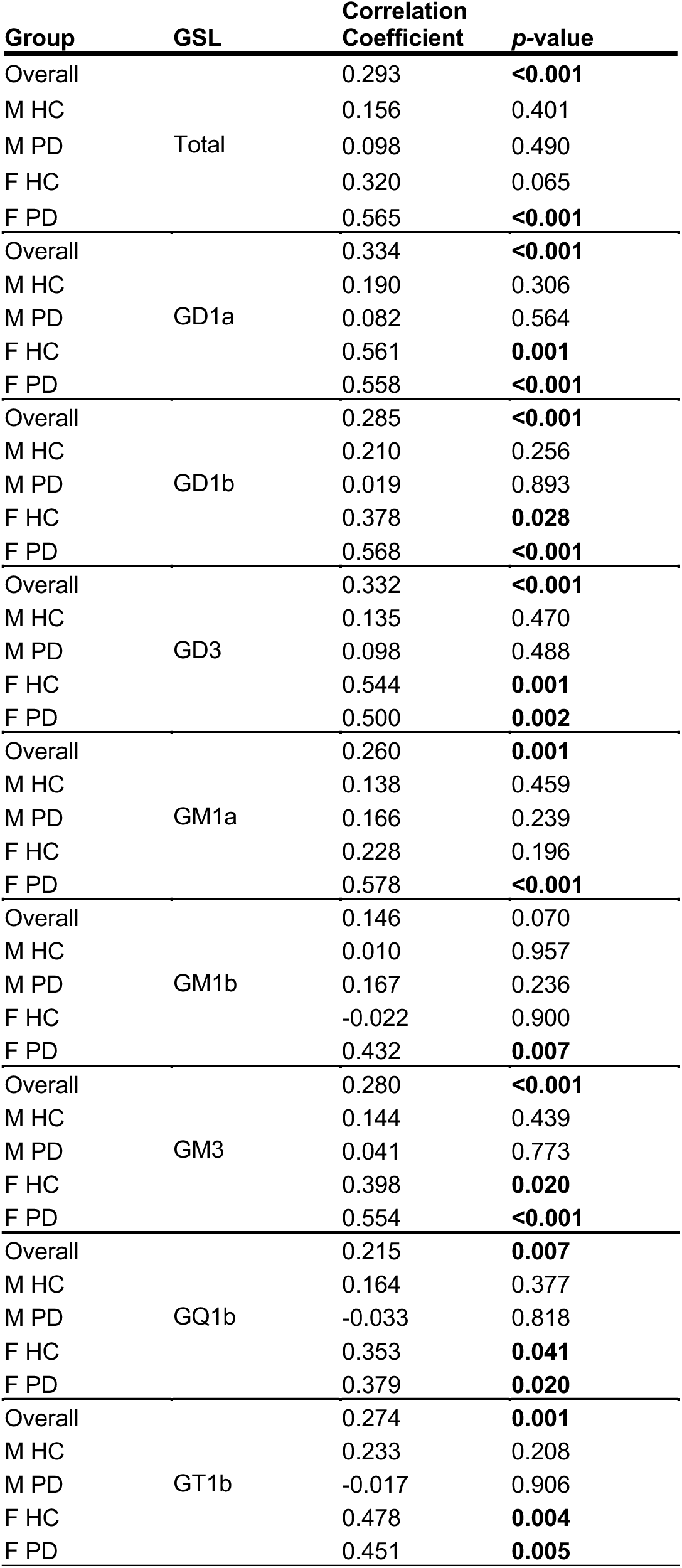
CSF GSLs positively correlate with CSF GPNMB levels in females but not males. Spearman-rank correlation analysis between major CSF gangliosides and CSF GPNMB levels (*n* = 157). Sub-group analysis stratified by disease state and sex; Male Healthy Controls (M HC, *n* = 31), Male Parkinson’s disease (M PD, *n* = 54), Female Healthy Controls (F HC, *n* = 34), Female Parkinson’s disease (F PD, *n* = 38).

Similarly, we analyzed the correlation between GSLs and GPNMB in plasma stratified by sex and disease status (**Table 2**). Again, GSLs in the female PD group correlated with most of the GSLs analyzed (Gb3, Gb4, GM1a, GM3 and total GSL) the only exception being GM2 that correlated with GPNMB only in the male control group. Gb3 and Gb4 correlated with GPNMB in the male PD group. We found no correlation between GPNMB and GSLs in the plasma of female controls (**Table 2**).

**Table 2:** Correlation between plasma GSLs and GPNMB levels. Sub-group analysis stratified by disease state and sex; Male Healthy Controls (M HC, n = 44), Male Parkinson’s disease (M PD, n = 72), Female Healthy Controls (F HC, n = 44), Female Parkinson’s disease (F PD, n = 44).

| Group | GSL | Correlation Coefficient | p-value |
| --- | --- | --- | --- |
| Overall |  | 0.198 | **0.005** |
| M HC | Total | 0.141 | 0.362 |
| M PD |  | 0.138 | 0.246 |
| F HC |  | -0.213 | 0.164 |
| F PD |  | 0.431 | **0.003** |
| Overall |  | 0.282 | **<0.001** |
| M HC | Gb3 | 0.215 | 0.161 |
| M PD |  | 0.282 | **0.016** |
| F HC |  | -0.046 | 0.766 |
| F PD |  | 0.417 | **0.005** |
| Overall |  | 0.29 | **<0.001** |
| M HC | Gb4 | 0.222 | 0.147 |
| M PD |  | 0.303 | **0.010** |
| F HC |  | 0.009 | 0.954 |
| F PD |  | 0.443 | **0.003** |
| Overall |  | 0.209 | **0.003** |
| M HC | GM1a | 0.197 | 0.200 |
| M PD |  | 0.123 | 0.305 |
| F HC |  | -0.099 | 0.521 |
| F PD |  | 0.475 | **0.001** |
| Overall |  | 0.227 | **0.001** |
| M HC | GM2 | 0.447 | **0.002** |
| M PD |  | 0.101 | 0.396 |
| F HC |  | -0.078 | 0.616 |
| F PD |  | 0.242 | 0.113 |
| Overall |  | 0.184 | **0.008** |
| M HC | GM3 | 0.117 | 0.450 |
| M PD |  | 0.179 | 0.133 |
| F HC |  | -0.163 | 0.289 |
| F PD |  | 0.36 | **0.016** |

### GPNMB levels correlated with age and α-synuclein levels in CSF

To further evaluate potential links between GPNMB levels in the CSF and clinical parameters, correlations between levels of GPNMB and other CSF biomarkers were determined. While GPNMB levels in the CSF and plasma did not correlate, GPNMB positively correlated with both age (**Figure 5A**) and α-synuclein levels in the CSF (**Figure 5B**). In contrast, in the plasma, GPNMB and α-synuclein levels were not significantly correlated (data not shown).

**Figure 5.**
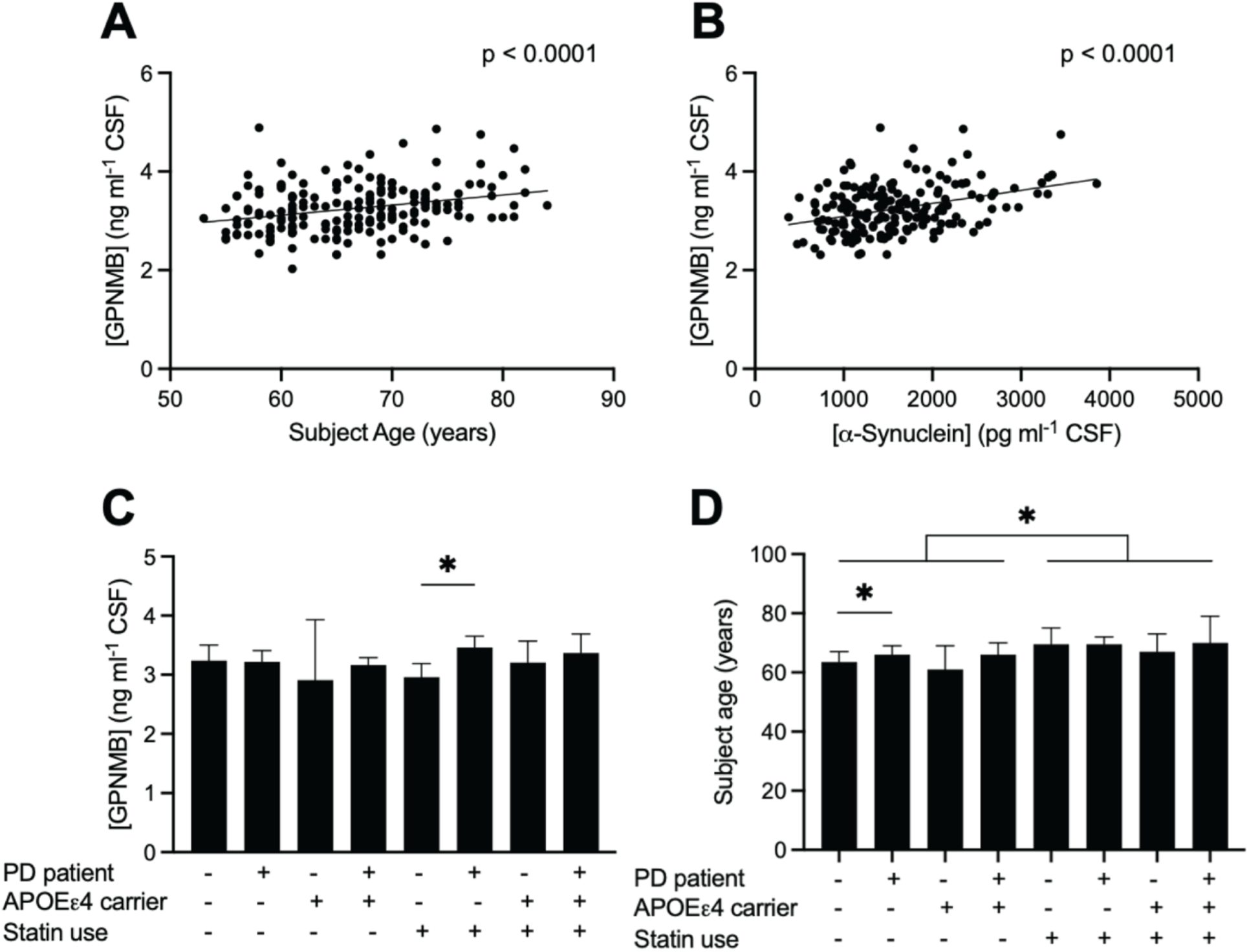
GPNMB levels correlated with age and α-synuclein levels in CSF. Scatter plots and simple linear regression of (**A**) subject age and GPNMB protein levels in the CSF (R² = 0.081), (**B**) α-synuclein and GPNMB protein levels in the CSF (R² = 0.131). The *p-*values represent the significance of the slopes. (**C**) GPNMB protein levels and (**D**) age in the CSF of control subjects (-) and PD patients (+) without or carrying the APOEε4 genotype and with or without concomitant statin medication. Median ± 95% confidence interval. Kruskal-Wallis test for main effects, Wilcoxon-Mann-Whitney tests for paired analyses adjusted by Holm-Sidak adjustment for multiple comparisons, \**p* < 0.05.

Concomitant disease processes and medications are common in older subjects. Given that the average age of the subjects was 67.1 ± 6.88 years (mean ± SD), molecular signatures of multimorbidity disease processes were likely to be present in the BioFind data set. However, the influence of pharmacological interventions on the CSF and plasma biochemistry of PD was unclear. The study subjects reported the concomitant use of medications from a list of 270 different brands that were analyzed as 14 drug classes (drug classes reported by more than 12 subjects were included for analysis of influences on clinical and biochemical data, supplements were excluded from analysis). Similar frequencies of drugs and drug combinations were reported by both control subjects and PD patients (**Supplementary Figure 6**).

Given previous findings that GPNMB levels correlate positively with triglyceride levels in PD substantia nigra ^9^, the subjects were stratified into PD and control groups based on the presence of at least one APOEε4 allele and concomitant use of medications that target cholesterol levels. The class of statin/cholesterol uptake inhibitors was comprised of Atorvastatin, Crestor, Ezetimibe with Simvastatin, Lipitor, Lovastatin, Pravachol, Pravastatin, Rosuvastatin, Simvastatin, Vytorin, Zocor and Zetia. In this analysis, the CSF of PD patients without an APOEε4 allele and using statins contained a 17% greater concentration of GPNMB than similar control subjects (**Figure 5C**). Indeed, statins were the only class of drugs that the subjects used concomitantly with PD drugs that linked GPNMB levels in the CSF to a diagnosis of PD. However, the use of concomitant medications can be determined by other factors. As an example, subjects using statins to control cholesterol levels were slightly older in the BioFind cohort (**Figure 5D**). Importantly, age did not explain the increased GPNMB levels in PD patients using statins but not carrying an APOEε4 allele. These data indicate that, among APOEε4 non-carriers, statin use is associated with higher CSF GPNMB levels in PD compared with controls, independent of age.

### GPNMB levels were associated with rs199347 genotype in plasma and CSF

The rs199347 single nucleotide polymorphism has been previously linked to PD risk and is associated with altered GPNMB expression in the brain ^27^. Analysis of GPNMB levels by rs199347 genotype revealed significant differences in both plasma (**Figure 6A**) and CSF (**Figure 6B**), indicating that GPNMB protein levels in both compartments are significantly associated with rs199347 genotype. In plasma, individuals carrying the A allele (GA and AA genotypes; representing the risk-associated variant) exhibited significantly higher GPNMB concentrations compared to those with the GG genotype (**Figure 6A**). A similar pattern was observed in CSF, where A allele carriers had significantly elevated GPNMB levels, with the highest levels detected in AA homozygotes (**Figure 6B**). When stratified by diagnosis, genotype-associated differences in GPNMB levels remained evident in both control and PD groups. Two-way ANOVA confirmed significant main effects of both genotype and diagnosis on GPNMB levels in plasma and CSF (**Figures 6C-D**).

**Figure 6.**
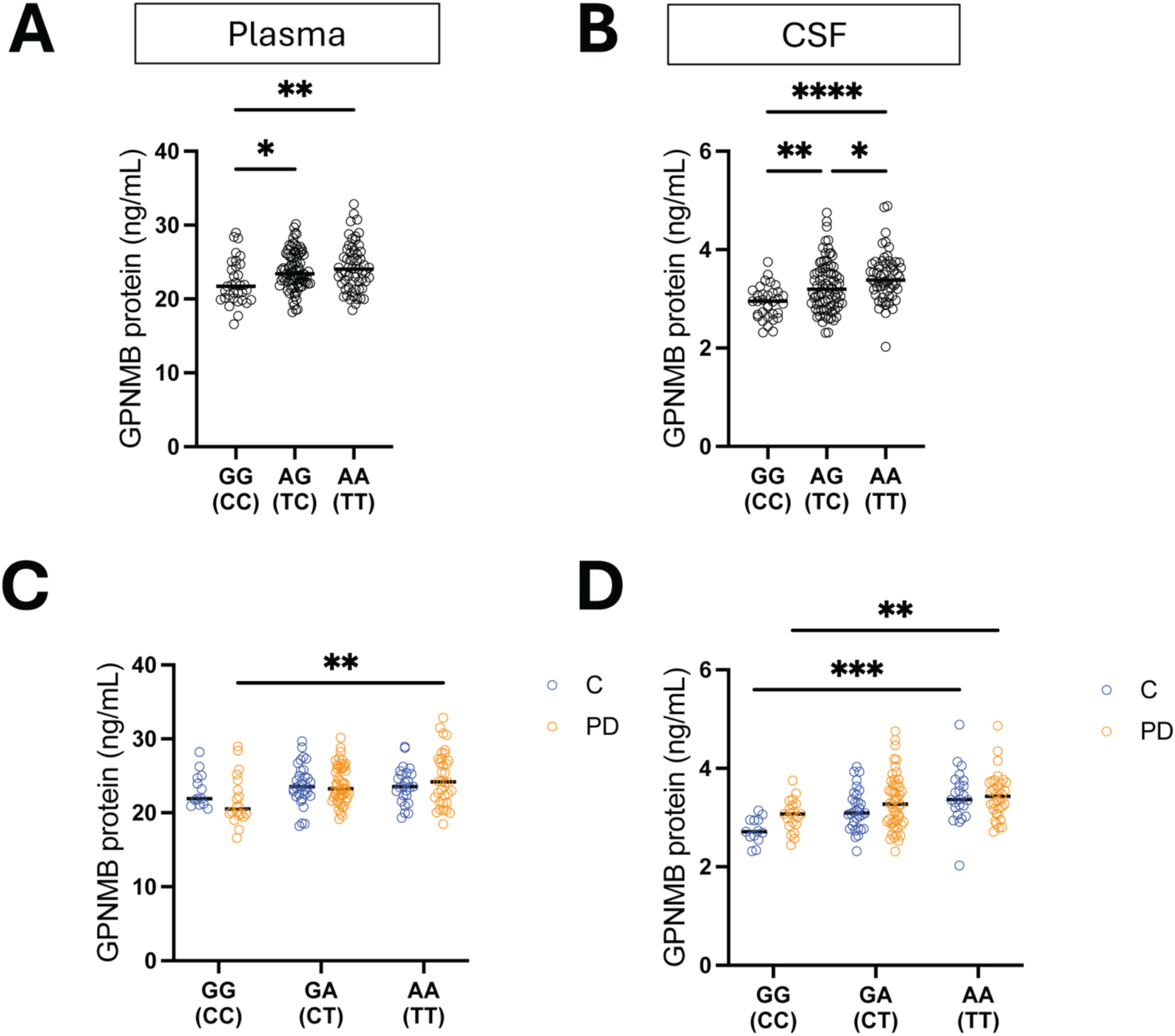
Biofluid GPNMB levels associate with the rs199347 genotype. **A)** Plasma and **B)** CSF GPNMB levels grouped by rs199347 genotype. G-to-A change on the forward DNA strand corresponds to a C-to-T change on the complementary strand. \**p*<0.05, \*\**p*<0.01, \*\*\**p*<0.001, \*\*\*\**p*<0.0001 Kruskal-Wallis test followed by Dunn’s adjustment for multiple comparisons. **C)** Plasma (*n* = 70 controls and 104 PD) and **D)** CSF (*n* = 68 controls and 105 PD) GPNMB levels grouped by rs199347 genotype and stratified by diagnosis. \*\**p* < 0.01, \*\*\**p* < 0.001 Two-way ANOVA followed by Tukey’s adjustment for multiple comparisons.

## Discussion

In this study, we have analyzed levels of GSLs in plasma and CSF from the BioFIND cohort. The HPLC technique we have used measures all GlcCer derived GSL species, allowing us to determine changes in individual species and total GSLs in a single assay. In plasma 3 out of the 14 GSLs measured changed significantly in PD relative to controls. Specifically, two paraglobosides were significantly increased in PD (alpha-2,3SpG and pGb) and GD1a + Le^b^ was significantly decreased in PD. It is interesting to note that Gb4, another globoside, has also been reported to be elevated in an independent cohort ^28^. What was also observed in the current study was that when these significant GSLs changes were stratified by sex, females had significantly higher levels of these specific GSLs than males. We therefore stratified all GSLs measured in plasma by sex. We found that all GSLs except LacCer and GlcCer were significantly higher in females relative to males independent of disease status. These results agree with a previous report comparing plasma from healthy male and female individuals ^29^. One hypothesis arising from these findings is that males are predisposed to PD relative to females in part because they have lower basal level of GSLs in plasma. For example, total gangliosides in plasma are significantly reduced in males relative to females, with a trend towards further reduction in the male PD group. Taken together these data could suggest that sex-related changes in GSL metabolism contribute to the risk of PD in males. It is interesting to note that of the three GSLs that were significantly different in PD versus control, in PD females, levels changed in the same direction as the males. This contrasts with the other GSLs that did not significantly change in PD, where control and PD females both had the same elevated levels relative to males. Further studies on larger cohorts of PD females will be required to determine whether levels of GSLs are linked to PD risk in females. The study cohort only included six *GBA1* carriers for plasma GSLs and GPNMB, and five and seven for CSF GSLs and GPNMB, respectively. No clear conclusion could be drawn from their distribution due to their underrepresentation in this cohort.

In contrast to the elevation in GSL species in the globo/paraglobo pathways (**Figure 3**), most gangliosides trended down in PD males relative to control males, in agreement with previous studies examining male dominated cohorts ^8, 28^. Although changes in individual GSLs did not always reach statistical significance, the trends mapped onto branches of the GSL biosynthetic pathway, suggesting a perturbation of specific pathways (**Supplementary Figure 7**).

The four major GSLs in CSF are the gangliosides GM1a, GD1a, GD1b and GT1b. It was interesting to note that the levels of all these gangliosides trended down in PD, again most prominently in males (**Supplementary Figure 5**). These gangliosides reside in the plasma membrane of cells of the CNS. Due to their biochemical properties, in CSF gangliosides are not free in solution but rather in complex with lipoproteins or as part of exosomes. The relevance of GSLs in CSF remains unclear as high levels of GSLs may reflect reduced levels of GSLs in the plasma membrane of cells of the CNS, potentially reducing neuroprotective gangliosides levels in neurons and glia. However, in this study these changes in GSLs in CSF did not reach statistical significance in relation to sex or PD status.

In addition to GSLs, we measured GPNMB in the same samples. There were no significant differences in GPNMB levels between PD and controls but as with GSLs there was a statistically significant sex effect, with females having higher levels of GPNMB in plasma and lower levels in CSF. Interestingly, the BioFIND dataset revealed not only correlations between GPNMB and GSLs but also a correlation of GPNMB with other PD relevant molecules, such as α-synuclein. These results suggest a broader network of lipid dysregulation and biomarker interconnectivity in PD in which the biological role of sex merits further characterization.

These findings have significant implications for PD biomarker research. They highlight the necessity of sex-specific analyses and the establishment of distinct biomarker reference ranges to account for sex differences. Such an approach will improve the precision of risk assessment and diagnostic criteria for PD. Furthermore, these insights are critical for future clinical trial design, emphasizing the need to stratify participants by sex to capture nuanced differences in biochemical markers and disease mechanisms.

### Anabolic and Catabolic Regulation of GSLs in the Context of Parkinson’s Disease Risk

The composition of GSLs is complex and is influenced by multiple factors, including flux through biosynthetic pathways, vesicular trafficking and lysosomal catabolism.

If male risk for PD is the net result of an altered pattern of GSLs or changes in charged to neutral GSLs in neuronal membranes, then those changes could arise through multiple independent mechanisms involving the contribution of multiple genes. Prior studies such as Robak *et al.* ^30^ underscored that PD pathophysiology may arise from perturbations beyond simple metabolic flux. This is further supported by diverse omics approaches uncovering genes implicated in PD in both anabolic (biosynthetic) and catabolic (lysosomal degradative) processes, which could underlie the observed shifts ^2, 4, 31^, along with genes involved in endo-lysosomal trafficking machinery ^32, 33^ which also influence GSL turnover. In line with broader observations of lysosomal dysfunction in PD, these diverse mechanisms could independently converge to disrupt GSL metabolism.

In addition, our sex-stratified analyses hint at a possible explanation for the male bias in PD incidence: males exhibited overall lower GSL levels in plasma, while females, particularly PD females, showed stronger correlations between GPNMB and GSLs. Interestingly, in CSF, GPNMB positively correlated with gangliosides in females irrespective of PD status. However, GM1 ganglioside only showed a significant correlation in PD females. As GM1 ganglioside has been shown to be neuroprotective in preclinical PD models ^34^ this relationship merits further investigation. This finding suggests that females who develop PD may carry distinct glycosphingolipid imbalances that could underlie critical protective or pathogenic mechanisms. Overall, the data argue for a broader view of PD risk where differential regulation of sphingolipid metabolism, both anabolic and catabolic, is influenced by age and sex.

### Age-associated changes in GPNMB in PD subjects

Aging is the most significant risk factor for PD and prior studies have established that GSL metabolism is significantly altered with age, particularly in the substantia nigra. Specifically, age-dependent reductions in GCase activity and complex gangliosides (including GD1a, GD1b, and GT1b) have been reported in both PD and normal aging, with corresponding accumulations of GlcCer and LacCer ^7, 8, 10^. Additionally, previous work has shown that in idiopathic PD fibroblasts, reduced GCase activity significantly correlates with decreased levels of LIMP2, the lysosomal trafficking receptor for GCase, suggesting that aging-related declines in lysosomal function may also involve impaired enzyme transport ^35^.

Our data demonstrate that CSF GPNMB levels increase with age in PD subjects and may reflect brain GSL elevations, such as GlcCer and its deacylated product GlcSph, which, although neither was measured in CSF in this study, have previously been shown to progressively increase with age in the brain in mice ^7^ and in the substantia nigra of PD patients ^8, 10^ . Notably, the relationship between GPNMB and age-dependent GSL changes suggests a broader dysregulation of GSL metabolism in PD, potentially linking ganglioside depletion, lysosomal impairment, and neuroinflammation in an age-dependent manner. Although no GSL changes associated with age were observed in this cohort, this highlights the need for longitudinal studies as the BioFind cohort lacks this dimension.

### Association of rs199347 genotype with increased GPNMB levels in PD and control subjects

The current findings demonstrate that the rs199347 SNP, located in a regulatory region near to the GPNMB gene, is significantly associated with elevated GPNMB protein levels in both CSF and plasma, independent of PD status. Individuals carrying the A allele, previously identified as a PD risk-associated variant ^36, 37^, consistently showed higher GPNMB concentrations, with the strongest effect observed in AA homozygotes. This confirms and extends earlier observations that rs199347 acts as an expression quantitative trait locus (eQTL) for GPNMB and is associated with increased GPNMB protein expression. Importantly, unlike prior studies, which observed rs199347-associated increases in GPNMB protein levels primarily in PD patients ^38^, the current findings demonstrate that this association extends to control subjects, suggesting that the SNP acts as a constitutive regulator of GPNMB protein levels in human biofluids, independent of disease status.

## Conclusions

This study shows that GSLs and GPNMB levels, in plasma and CSF, are closely associated in PD and significantly influenced by sex, age, and genetic background. Plasma GSL alterations in PD, together with lower overall GSL levels in males and stronger GPNMB-GSL correlations in females, suggest that sex-specific lipid dysregulation may contribute to differential susceptibility to PD. Furthermore, increased CSF GPNMB levels with age, correlation with α-synuclein, and association with the PD-risk rs199347 variant support GPNMB as a biomarker of lysosomal dysfunction and PD-related neurodegenerative processes.

## Final remarks

One of the challenges of investigating clinical cohorts is that they can vary in several ways. This includes patient numbers, genetic background, sex balance, stage of disease and concomitant use of medications and smoking history. In the future, it will be important to control better for these factors with the goal of teasing out underlying mechanisms of risk. The analysis of BioFind has provided a single snapshot in time of a well-researched cohort. It offers the possibility of cross correlating these findings with other parameters/biomarkers that have been measured in the same set of samples. This cohort had the advantage of having a 60:40 sex composition (M:F), which has proved invaluable for unmasking the sex effects that we report here. Longitudinal analysis of PD and controls in the future would therefore be instrumental for better understanding the mechanisms underpinning PD risk.

## Acknowledgments

Data used in the preparation of this article were obtained from the Fox Investigation for New Discovery of Biomarkers (“BioFIND”) database (http://biofind.loni.usc.edu/). For up-to-date information on the study, visit https://www.michaeljfox.org/biospecimens. BioFIND is sponsored by The Michael J. Fox Foundation for Parkinson’s Research (MJFF) with support from the National Institute for Neurological Disorders and Stroke (NINDS). This study was funded by the Michael J Fox Foundation for Parkinson’s Research Award # 010388. Authors MFS, RB, DTV, DP, JH and FP were also funded by the Aligning Science Across Parkinson’s grant ASAP-000478.

## Author Contributions

PH, OI, JH, and FP contributed to the conception and design of the study and supervision; MFS, EDB, RB, DTV, DP, MCB, and OC, contributed to the acquisition and analysis of data; MFS, EDB, RB, MCB, OC, PH, OI, and FP contributed to drafting the text or preparing the figures.

## Data Availability

The data generated in this study are available on the BioFIND website (https://biofind.loni.usc.edu/download-data.php), study 143. An application is required to access the data. The code, protocols, and key lab materials used and generated in this study are listed in a Key Resources Table alongside their persistent identifiers at DOI: 10.5281/zenodo.20094251. An earlier version of this manuscript was posted to BioRxiv on 13 April 2026 at https://doi.org/10.64898/2026.04.09.712000.

## References

1. Cooper O, Hallett P, Isacson O. Upstream lipid and metabolic systems are potential causes of Alzheimer’s disease, Parkinson’s disease and dementias. Febs j. 2024 Feb;291(4):632–45.

2. Hallett PJ, Engelender S, Isacson O. Lipid and immune abnormalities causing age-dependent neurodegeneration and Parkinson’s disease. J Neuroinflammation. 2019 Jul 22;16(1):153.

3. Isacson O, Brekk OR, Hallett PJ. Novel Results and Concepts Emerging From Lipid Cell Biology Relevant to Degenerative Brain Aging and Disease. Frontiers in neurology. 2019;10:1053.

4. Wallom KL, Fernández-Suárez ME, Priestman DA, et al. Glycosphingolipid metabolism and its role in ageing and Parkinson’s disease. Glycoconj J. 2021 Nov 10.

5. Dehay B, Bové J, Rodríguez-Muela N, et al. Pathogenic lysosomal depletion in Parkinson’s disease. J Neurosci. 2010 Sep 15;30(37):12535–44.

6. Pascua-Maestro R, Diez-Hermano S, Lillo C, Ganfornina MD, Sanchez D. Protecting cells by protecting their vulnerable lysosomes: Identification of a new mechanism for preserving lysosomal functional integrity upon oxidative stress. PLoS genetics. 2017 Feb;13(2):e1006603.

7. Hallett PJ, Huebecker M, Brekk OR, et al. Glycosphingolipid levels and glucocerebrosidase activity are altered in normal aging of the mouse brain. Neurobiol Aging. 2018 Jul;67:189–200.

8. Huebecker M, Moloney EB, van der Spoel AC, et al. Reduced sphingolipid hydrolase activities, substrate accumulation and ganglioside decline in Parkinson’s disease. Mol Neurodegener. 2019 Nov 8;14(1):40.

9. Brekk OR, Honey JR, Lee S, Hallett PJ, Isacson O. Cell type-specific lipid storage changes in Parkinson’s disease patient brains are recapitulated by experimental glycolipid disturbance. Proc Natl Acad Sci U S A. 2020 Nov 3;117(44):27646–54.

10. Rocha EM, Smith GA, Park E, et al. Progressive decline of glucocerebrosidase in aging and Parkinson’s disease. Ann Clin Transl Neurol. 2015 Apr;2(4):433–8.

11. Ripoll VM, Irvine KM, Ravasi T, Sweet MJ, Hume DA. Gpnmb is induced in macrophages by IFN-gamma and lipopolysaccharide and acts as a feedback regulator of proinflammatory responses. J Immunol. 2007 May 15;178(10):6557–66.

12. Marques AR, Gabriel TL, Aten J, et al. Gpnmb Is a Potential Marker for the Visceral Pathology in Niemann-Pick Type C Disease. PLoS One. 2016;11(1):e0147208.

13. Murugesan V, Liu J, Yang R, et al. Validating glycoprotein non-metastatic melanoma B (gpNMB, osteoactivin), a new biomarker of Gaucher disease. Blood cells, molecules & diseases. 2018 Feb;68:47–53.

14. Zigdon H, Savidor A, Levin Y, Meshcheriakova A, Schiffmann R, Futerman AH. Identification of a biomarker in cerebrospinal fluid for neuronopathic forms of Gaucher disease. PLoS One. 2015;10(3):e0120194.

15. Kramer G, Wegdam W, Donker-Koopman W, et al. Elevation of glycoprotein nonmetastatic melanoma protein B in type 1 Gaucher disease patients and mouse models. FEBS open bio. 2016 Sep;6(9):902–13.

16. Vardi A, Zigdon H, Meshcheriakova A, et al. Delineating pathological pathways in a chemically induced mouse model of Gaucher disease. The Journal of pathology. 2016 Aug;239(4):496–509.

17. Kumaran R, Cookson MR. Pathways to Parkinsonism Redux: convergent pathobiological mechanisms in genetics of Parkinson’s disease. Hum Mol Genet. 2015 Oct 15;24(R1):R32–44.

18. Neal ML, Boyle AM, Budge KM, Safadi FF, Richardson JR. The glycoprotein GPNMB attenuates astrocyte inflammatory responses through the CD44 receptor. J Neuroinflammation. 2018 Mar 8;15(1):73.

19. Bogacki EC, Longmore G, Lewis PA, Herbst S. GPNMB is a biomarker for lysosomal dysfunction and is secreted via LRRK2-modulated lysosomal exocytosis. bioRxiv. 2025:2025.01.01.630988.

20. Moloney EB, Moskites A, Ferrari EJ, Isacson O, Hallett PJ. The glycoprotein GPNMB is selectively elevated in the substantia nigra of Parkinson’s disease patients and increases after lysosomal stress. Neurobiol Dis. 2018 Dec;120:1–11.

21. Kang UJ, Goldman JG, Alcalay RN, et al. The BioFIND study: Characteristics of a clinically typical Parkinson’s disease biomarker cohort. Mov Disord. 2016 Jun;31(6):924–32.

22. Neville DC, Coquard V, Priestman DA, et al. Analysis of fluorescently labeled glycosphingolipid-derived oligosaccharides following ceramide glycanase digestion and anthranilic acid labeling. Anal Biochem. 2004 Aug 15;331(2):275–82.

23. Priestman DA, te Vruchte D, Wallom KL, et al. Analysis of glycosphingolipids from human plasma. protocolsio. 2021.

24. Priestman DA, Bush R, Leondaraki M, et al. Analysis of glycosphingolipids from human cerebrospinal fluid V.1 protocolsio. 2023.

25. Goldman JG, Andrews H, Amara A, et al. Cerebrospinal fluid, plasma, and saliva in the BioFIND study: Relationships among biomarkers and Parkinson’s disease Features. Mov Disord. 2018 Feb;33(2):282–8.

26. Haaxma CA, Bloem BR, Borm GF, et al. Gender differences in Parkinson’s disease. J Neurol Neurosurg Psychiatry. 2007 Aug;78(8):819–24.

27. Murthy MN, Blauwendraat C, Guelfi S, Hardy J, Lewis PA, Trabzuni D. Increased brain expression of GPNMB is associated with genome wide significant risk for Parkinson’s disease on chromosome 7p15.3. Neurogenetics. 2017 Jul;18(3):121–33.

28. Te Vruchte D, Sturchio A, Priestman DA, et al. Glycosphingolipid Changes in Plasma in Parkinson’s Disease Independent of Glucosylceramide Levels. Mov Disord. 2022 Oct;37(10):2129–34.

29. Sales S, Graessler J, Ciucci S, et al. Gender, Contraceptives and Individual Metabolic Predisposition Shape a Healthy Plasma Lipidome. Sci Rep. 2016 Jun 14;6:27710.

30. Robak LA, Jansen IE, van Rooij J, et al. Excessive burden of lysosomal storage disorder gene variants in Parkinson’s disease. Brain. 2017 Dec 1;140(12):3191–203.

31. Wang H, Zhao M, Chen G, Lin Y, Kang D, Yu L. Identifying MSMO1, ELOVL6, AACS, and CERS2 related to lipid metabolism as biomarkers of Parkinson’s disease. Sci Rep. 2024 Jul 30;14(1):17478.

32. Schormair B, Kemlink D, Mollenhauer B, et al. Diagnostic exome sequencing in early-onset Parkinson’s disease confirms VPS13C as a rare cause of autosomal-recessive Parkinson’s disease. Clin Genet. 2018 Mar;93(3):603–12.

33. Isacson O. Lysosomes to combat Parkinson’s disease. Nat Neurosci. 2015 May 26;18(6):792–3.

34. Schneider JS, Aras R, Williams CK, Koprich JB, Brotchie JM, Singh V. GM1 Ganglioside Modifies α-Synuclein Toxicity and is Neuroprotective in a Rat α-Synuclein Model of Parkinson’s Disease. Sci Rep. 2019 Jun 10;9(1):8362.

35. Thomas R, Moloney EB, Macbain ZK, Hallett PJ, Isacson O. Fibroblasts from idiopathic Parkinson’s disease exhibit deficiency of lysosomal glucocerebrosidase activity associated with reduced levels of the trafficking receptor LIMP2. Mol Brain. 2021 Jan 19;14(1):16.

36. Chang D, Nalls MA, Hallgrimsdottir IB, et al. A meta-analysis of genome-wide association studies identifies 17 new Parkinson’s disease risk loci. Nat Genet. 2017 Oct;49(10):1511–6.

37. Nalls MA, Blauwendraat C, Vallerga CL, et al. Identification of novel risk loci, causal insights, and heritable risk for Parkinson’s disease: a meta-analysis of genome-wide association studies. Lancet Neurol. 2019 Dec;18(12):1091–102.

38. Diaz-Ortiz ME, Seo Y, Posavi M, et al. GPNMB confers risk for Parkinson’s disease through interaction with α-synuclein. Science. 2022 Aug 19;377(6608):eabk0637.

