## Supplementary Data for "GPNMB and glycosphingolipid measurements in cerebrospinal fluid and plasma from Parkinson’s disease patients"

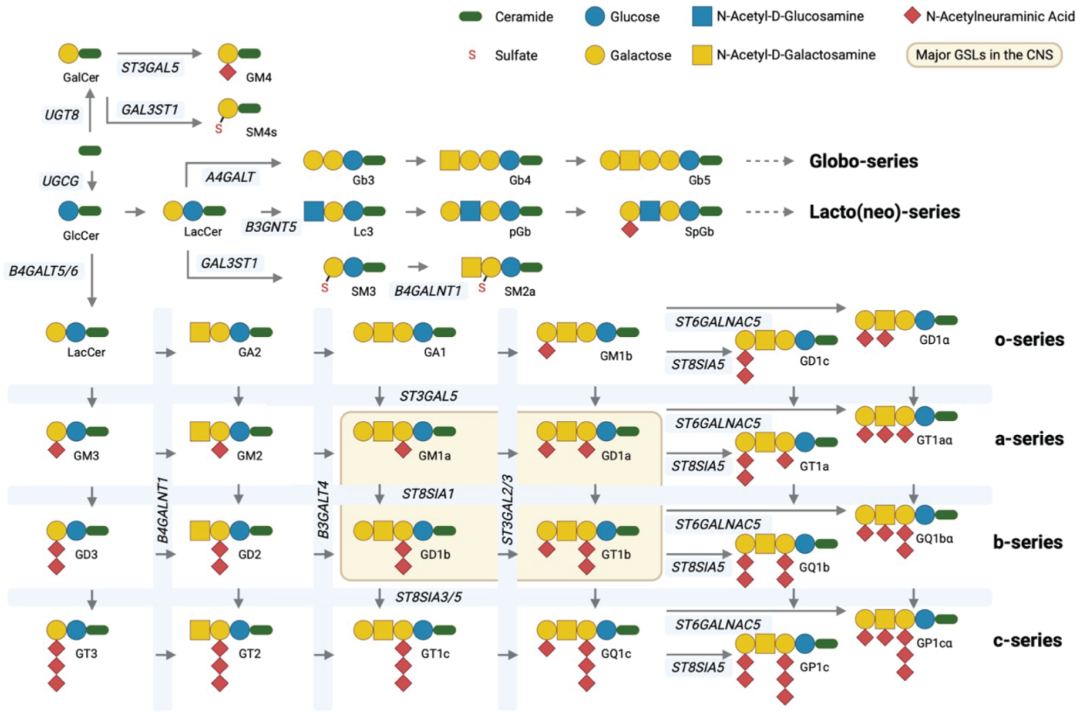


**Supplementary Figure 1. GSL Biosynthesis pathway.** Overview of the biosynthetic pathway of GSLs in humans, along with the genes encoding the enzymes responsible for their processing. Adapted from [23].

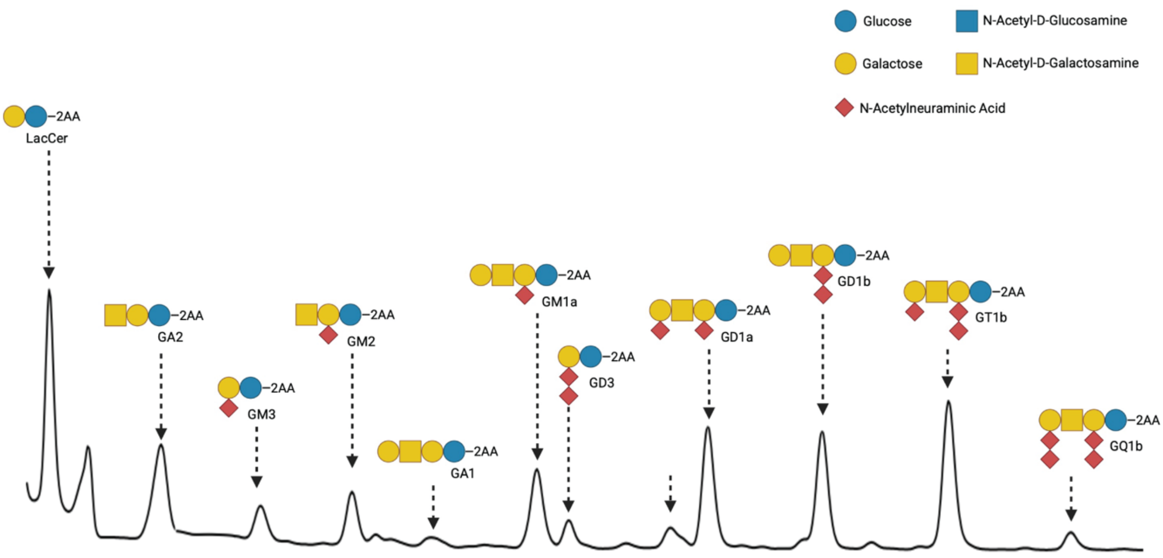


**Supplementary Figure 2. HPLC profile of CSF GSLs.** Typical HPLC profile of 2-AA labelled glycans enzymatically released from GSLs in human CSF. Known peaks are labelled and their glycan structures are shown. Adapted from [24].

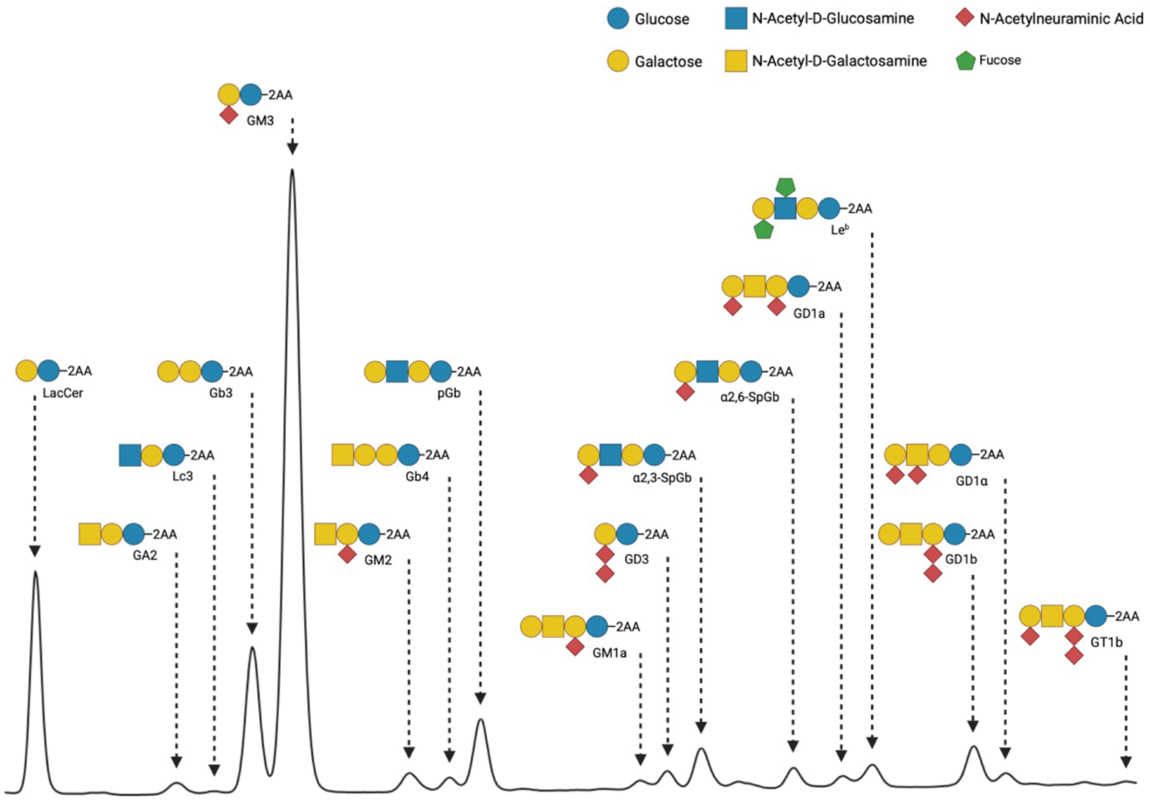


**Supplementary Figure 3. HPLC profile of plasma GSLs.** Typical HPLC profile of 2-AA labelled glycans enzymatically released from GSLs in human plasma. Known peaks are labelled and their glycan structures are shown. Adapted from [23].

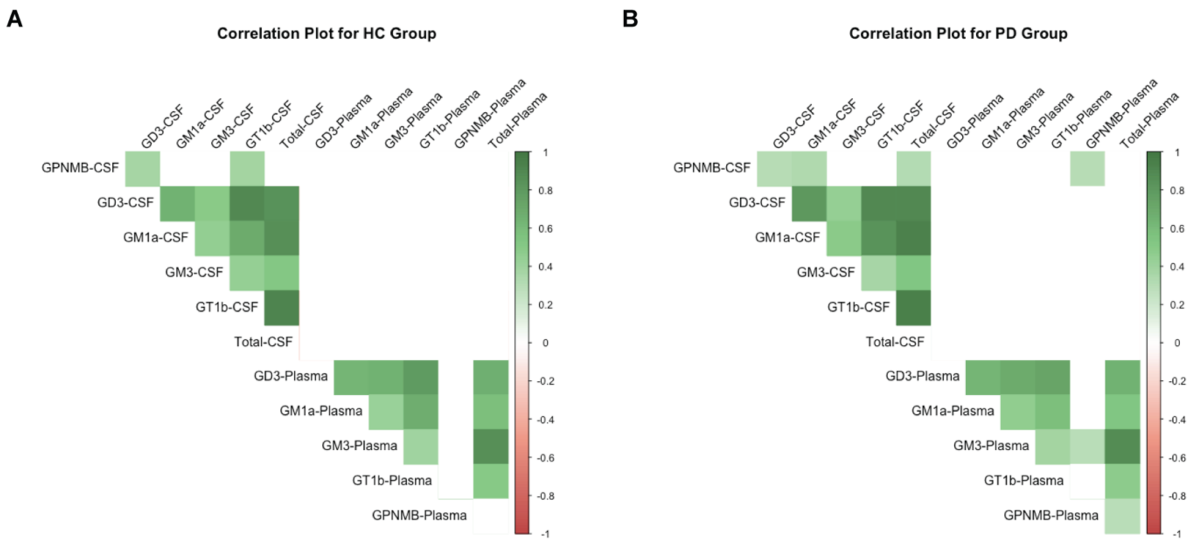


**Supplementary Figure 4: No correlation was found in individual GSL levels in plasma with those in CSF.** Spearman-rank correlations of CSF and plasma GSLs in healthy controls (A) or PD patients (B), visualized as correlograms. Shading of cells represents the magnitude of correlation coefficient, statistically non-significant correlations visualized as blank cells. Threshold for statistical significance was $p < 0.01$.

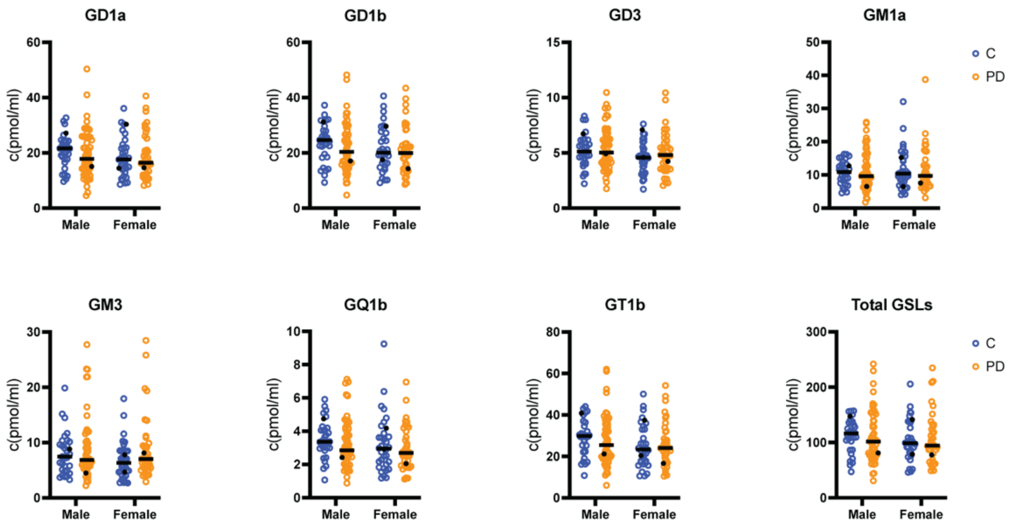


**Supplementary Figure 5. GSL levels in CSF.** Quantified individual GSL species from CSF (*n* = 157), stratified by sex (*n* = 85 male and 72 female) and disease status (*n* = 65 controls and 92 PD). Individual subjects visualized as open circles, with controls (blue), PD (orange), and *GBA1* mutant carriers as filled black circles (*n* = 5). The median for each group is indicated with a black line. One male control subject was excluded from the analysis as an outlier. Therefore, *n* = 156 allowing for the removal of this sample. Linear regression of log-normalized values of plasma GSL species showed no statistical difference between groups.

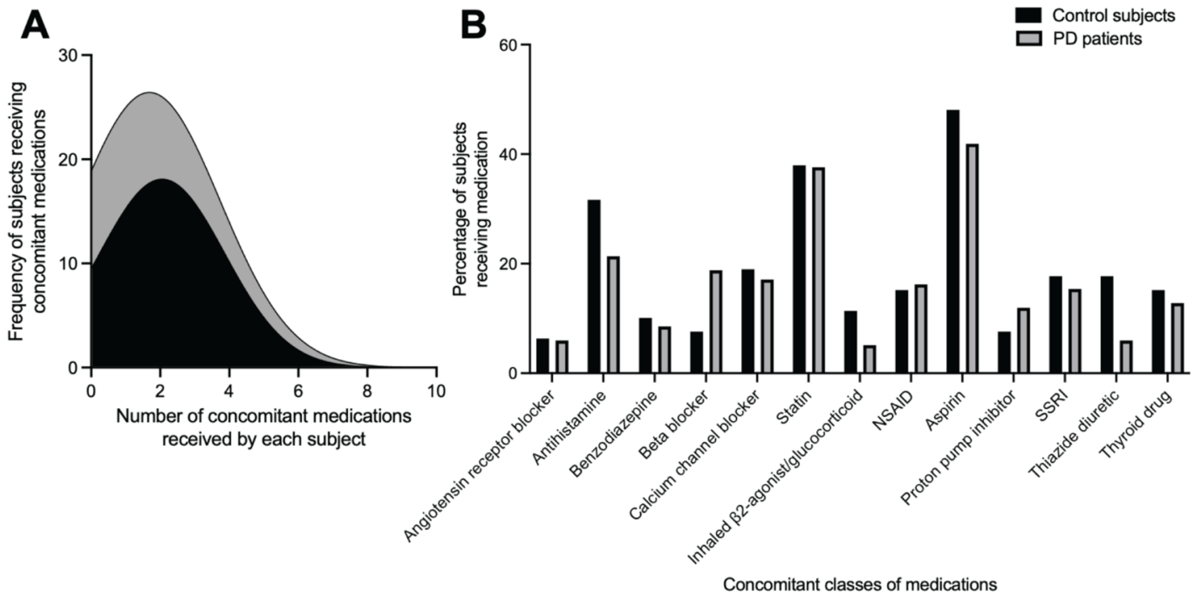


**Supplementary Figure 6. Control subjects and PD patients received similar combinations of concomitant medications.** (A) Frequency distributions of the number of concomitant medications received by either the control subjects (black) or PD patients (grey). (B) The percentage of control subjects (black bars) or PD patients (grey bars) that received different classes of medication.

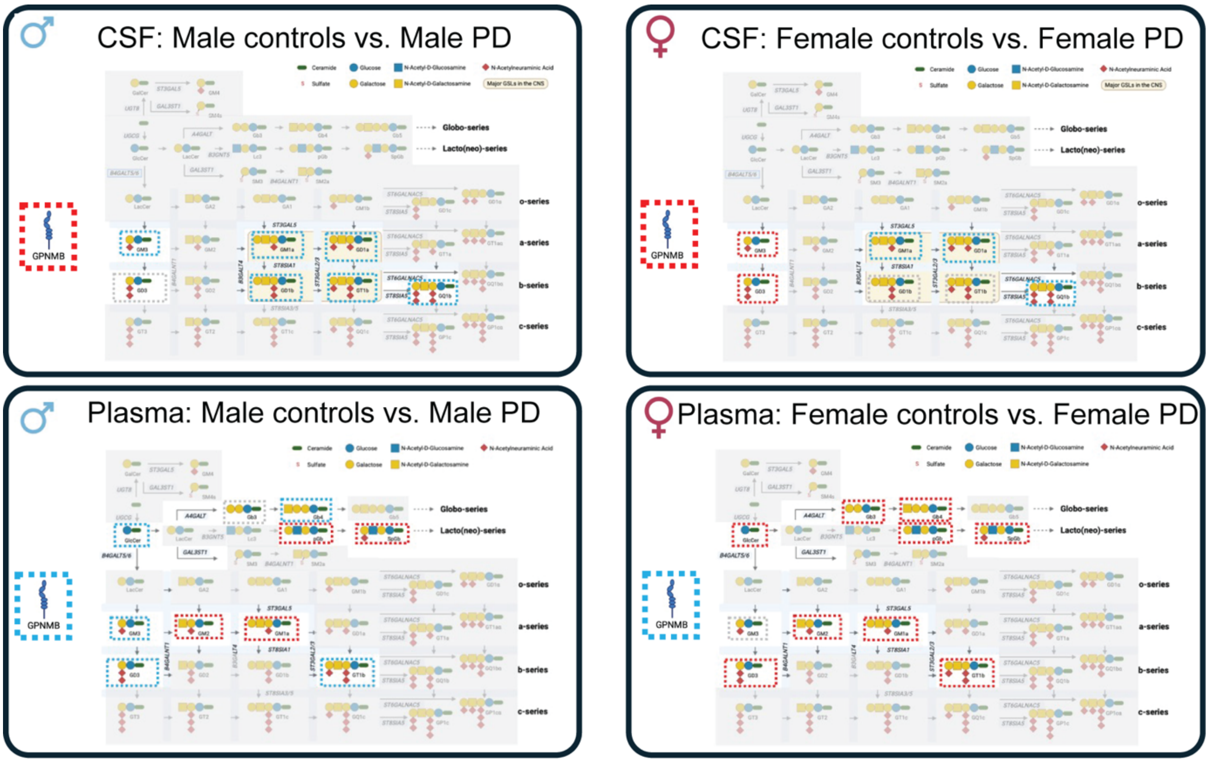


**Supplementary Figure 7. Observed changes in GSLs and GPNMB in the context of biosynthetic pathway.** The four panels depict trends of changes in glycosphingolipids (GSLs) and GPNMB levels in Parkinson's disease (PD) versus control subjects, stratified by biofluid and sex. Trends are indicated with dashed boxes surrounding the respective analytes: higher in PD (red), lower in PD (blue) or no trend (grey).

| Biofluid | Analytes | Diagnosis | Total | Male | Female |
|---|---|---|---|---|---|
| **CSF** | GSLs | Control | 65 | 31 | 34 |
| | | PD | 92 | 54 | 38 |
| | GPNMB | Control | 78 | 36 | 42 |
| | | PD | 107 | 64 | 43 |
| **Plasma** | GSLs | Control | 88 | 44 | 44 |
| | | PD | 116 | 72 | 44 |
| | GPNMB | Control | 89 | 45 | 44 |
| | | PD | 116 | 72 | 44 |

**Supplementary Table 1: Overview of sample numbers for CSF and plasma used in the analysis for this study.**
